# Imperfect gold standard gene sets yield inaccurate evaluation of causal gene identification methods

**DOI:** 10.1101/2023.05.04.539407

**Authors:** Lijia Wang, Xiaoquan Wen, Jean Morrison

## Abstract

Causal gene discovery methods are often evaluated using gold-standard (GS) sets of causal genes. However, GS gene sets are always incomplete, leading to mis-estimation of sensitivity, specificity, AUC. Labeling biases in GS gene sets can also lead to inaccurate ordering of discovery methods. We argue that evaluation of these methods should rely on statistical techniques like those used for variant discovery, rather than on comparison with GS gene sets.

---

Identifying causal genes for complex diseases can highlight disease-specific disregulated pathways, improve disease classification, and identify drug targets [3]. In genetic association analysis, it has become a common practice to implicate putative causal genes (PCG) computationally by linking variant-level genetic association evidence and the existing biological knowledge base [7]. Some of those computational methods approach PCG implication as a supervised learning problem, aiming to predict unknown binary causal/non-causal gene labels for a given trait. Others rank the candidate genes by their likelihood of being PCGs, returning a continuous probability estimate or ranking for each gene. In this article, we focus on the common practice of evaluating PCG implication methods in reference to known (gold standard, GS) sets of causal genes. A critical challenge for this assessment strategy is that all GS gene sets are incomplete and known causal genes may differ meaningfully from as-yet unidentified causal genes. We show that when the GS gene set is incomplete, estimates of power, specificity, receiver operating characteristic (ROC) will be inaccurate and may even misorder the relative quality of two different classifiers. This phenomenon can occur even if the GS gene set contains no erroneously labeled non-causal genes. We argue that no true GS sets of labeled genes are currently available and therefore urge caution in interpreting comparisons of causal gene classifiers made based on existing sets of labels.

Table 1 summarizes major GS gene set curation methods. We note that the labeled causal genes by these methods are typically reliable, as the standards for causality claim are usually stringent. However, some causal genes will inevitably be missed (thus motivating continued PCG discovery research). When the curated GS gene sets are used as reference sets for evaluation, genes not labeled as positive are implicitly treated as non-causal. However, the more confident we are in the positive labels of a GS gene set, the more mislabeled non-causal genes we can expect [7]. GS gene sets also tend to favor genes with particular features determined by the method of constructing the set. For example, GS gene sets derived from the set of causal coding variants favor genes that act through protein-coding changes rather than expression regulatory mechanisms [8]. Most classification methods also have gene-feature-related biases due to the type of data they use as input. A PCG implication method will appear more accurate if it is evaluated using a GS gene set with similar feature-related biases to its own and less accurate if the GS gene set has different biases. Authors naturally select a GS gene set constructed using features they feel are important and may, therefore, unintentionally tilt the scales toward their own proposed method.

**Table 1:** Constructions of Gold Standard gene sets. All of the listed methods used their constructed gold standard gene sets to compare the precision and recall of classification methods, or evaluate the AUC of ranking methods.

| Construction scheme | Publication |
| --- | --- |
| Genes harboring large-effect-size coding variants for selected phenotypes derived from OMIM and additional literature. | Connally et. al. (2021) [S1] |
| Genes that are targets of any known drug against the disease of interest (reported in Open Targets). | Picart-Armada et. al. (2019) [S7] |
| Genes annotated to phenotypes of interest in databases (OMIM, GO, DrugBank, literature derived drug targets) | Greene et. al. (2015) [S3] Tranchevent et. al. (2016) [S8] |
| Phenotype-associated genes compiled from GWAS catalog mining, publication-based, and expert curation. | Kolosov et. al (2021) [S5] <sup>1</sup> |
| Genes derived from expert domain knowledge, drug target-disease pairs, experimental alteration, and functional data. | Mountjoy et. al. (2021) [S6] |
| Genes with fine-mapped protein coding variants within loci non-coding credible sets. | Weeks et. al. (2020) [S9] |
| Top 10% genes ranked by PoPS for validation. | Gazal et. al. (2022) [S2] |
| Phenotype associated genes in recent reports from the Broad, TOPMed consortium, and IntOGen project. | Kessler et. al. (2021) [S4] <sup>2</sup> |

A more accurate view of genes outside the constructed GS set is not as negatives (N) but as unlabeled (U). Combined with accurately positively labeled genes in the GS set, the overall GS gene set should be regarded as positive-unlabeled (PU) data, a term used in semi-supervised machine learning. Using PU data to evaluate performance as though they were positive-negative (PN) labeled data results in inaccurate evaluations [4]. A PU-labeled gene set with perfect positive labeling consists of three subsets of genes: true causal genes that are correctly identified (labeled positives, LP), true causal genes that are not labeled and therefore assumed to be non-causal (unlabeled positives, UP), and non-causal genes that are unlabeled and therefore correctly assumed to be non-causal (unlabeled negatives, UN) (Fig. 1a). Evaluation treating PU labels (with perfect positive labeling) as PN labels will always underestimate the precision (or positive predictive value) and overestimate the negative predictive value (NPV) of a classifier (Fig. 1b). If we further assume that inaccuracies in the PU data are random with respect to features that influence classification accuracy, evaluation based on PU-labeled data will correctly estimate sensitivity and overestimate specificity. However, if LP and UP genes differ on important features, sensitivity and specificity may be either over- or under-estimated (Appendix A). In genetics research, we expect labeling biases because there are multiple molecular mechanisms by which a causal gene can affect complex diseases, and different classification and GS identification methods will favor different mechanisms. Error in estimating sensitivity and specificity results in error in the ROC curve and therefore error in the area under the ROC curve (AUC). This means that this error applies to evaluation of ranking methods as well as methods that return only hard classifications.

**Fig. 1:**
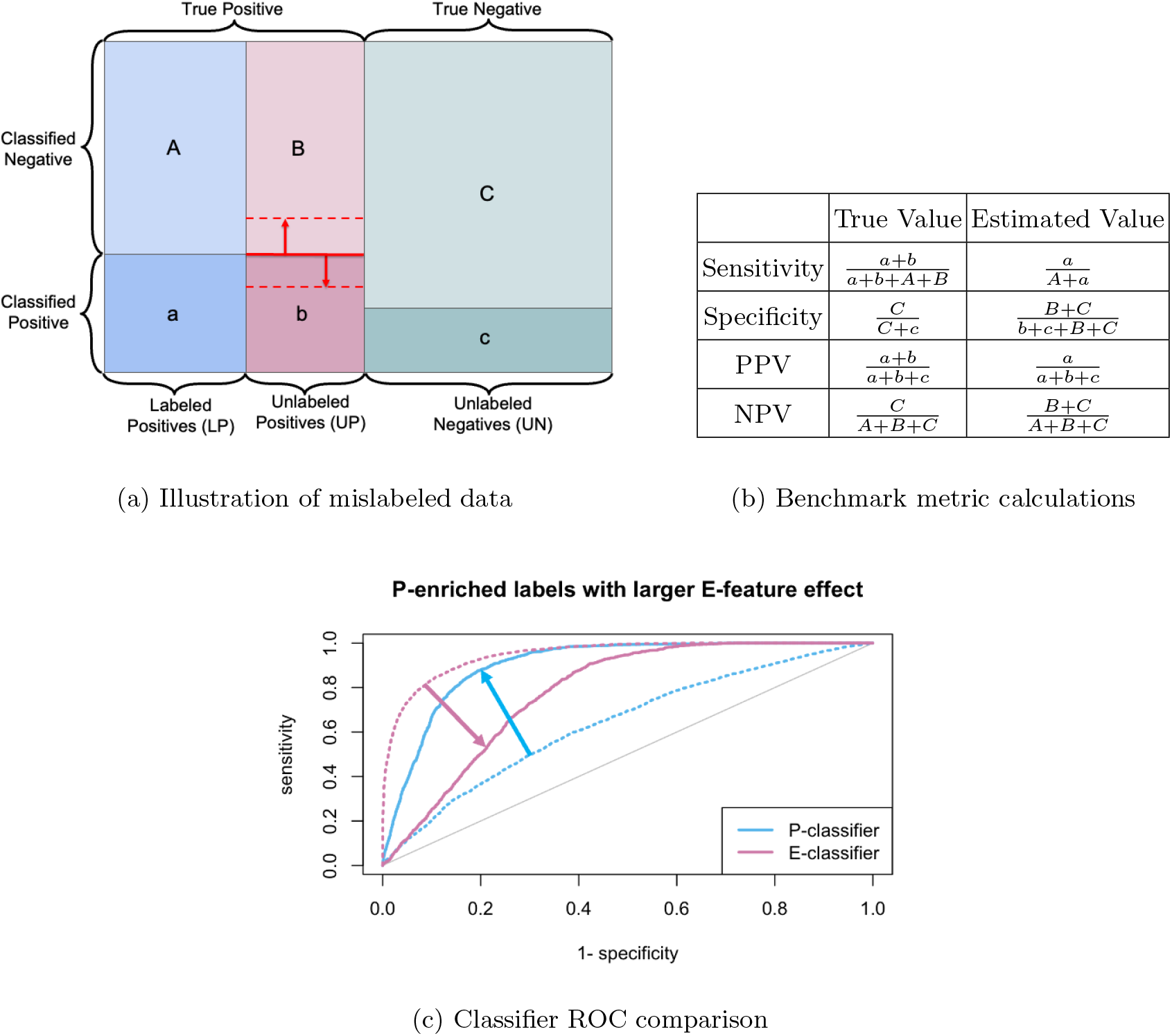
Comparison of Classifiers on True vs. Biased Labels. (a) Illustration of classifier performance relative to a typical GS gene set. The classifier may not have the same performance on UP genes as it has on LP genes (red lines and arrows), which affects estimated sensitivity and specificity. (b) Calculation for true vs. estimated accuracy in mislabeled data. PPV stands for positive predictive value and NPV stands for negative predictive value. (c) ROC curves for simulated classifiers. Dotted lines show true classifier performance, solid lines show performance estimated using an incomplete GS gene set. Arrows indicate the difference between true and estimated curves.

To illustrate this issue, we consider a hypothetical example in which each gene has two continuous, measurable features, *P* and *E*. We think of these features as continuous summaries of the evidence that a gene acts on the trait through mechanisms mediated by either protein sequence (*P*) or expression level (*E*). Causal genes with high values of *P* are more likely to be labeled positive in the GS gene set than those with high levels of *E* but low levels of *P*. We refer to these as P-enriched causal gene lables. We consider two hypothetical PCG classifiers. The E-Classifier, has higher sensitivity to detect causal genes with high *E* values and the P-Classifier has higher sensitivity to detect causal genes with high *P* values. Figure 1c shows that the ROC curve for the E-Classifier is below the truth, resulting in a downward bias in estimated AUC. In comparison, the evaluation for the P-Classifier, is overly optimistic. In this particular setting, the E-Classifier has a better overall ability to identify PCGs than the P-Classifier but is evaluated being much worse.

Due to the myriad biological mechanisms leading to complex phenotypes, it is currently impossible to confidently determine a comprehensive GS gene set that includes all causal genes for any trait. Several studies have acknowledged that supervised ML methods designed to classify PCGs should not be trained on incomprehensive GS gene sets [2], [5]. Here, we want to draw attention to the fact that sensitivity and specificity estimated using incomplete labels are also inaccurate, making it inappropriate to compare and evaluate methods using these measures or AUC estimated using GS gene sets. To address a similar issue in other fields, researchers have proposed incorporating negative controls (i.e., known negatives) or weights estimating each unit’s probability of being detected based on its features ([6], [1]). However, whether either approach is feasible for the PCG implication problem is unclear. An alternative approach that circumvents the issue is to use a statistical model-based approach for causal gene identification. Methods based on estimating parameters in probabilistic models provide model-based measures of uncertainty, such as posterior inclusion probabilities or confidence intervals. These methods can also be evaluated in simulations to test their sensitivity to violations of modeling assumptions. Such a paradigm has been established as a long-standing practice in genetics research to identify causal variants. Finally, we re-emphasize that researchers should avoid concluding the relative performance of different classifiers based on performance measures estimated using PU-labeled gene sets and should clearly acknowledge the limitations of any labeled gene set used in evaluation or training.

Simulation procedure and results can be found at https://lijiaw.gitlab.io/GS-gene-sets/comparison.html.

## 1 Appendix

### A Illustrating Effect of Mislabeling on Classification Accuracy

Receiver operating characteristics (ROC) analysis is one of the most common diagnostic methods for binary classifications, and is constructed using a combination of sensitivity and specificity estimations. In this section we will demonstrate how labeling some true positives as negatives in the reference gold standard data set affects estimates of sensitivity and specificity.

A set of binary, labeled data is illustrated in Figure A.1, where a portion of the true positives are labeled as negative. Definitions of sensitivity, specificity, negative predictive value (NPV) and positive predictive value (PPV) are given in Table 1b. From these definitions, we can establish that, in all cases,

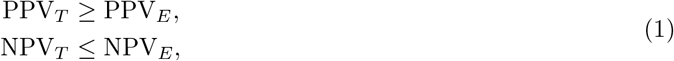

where subscripts *T* and *E* refer to the true and estimated values respectively.

Figure A.1 shows four possible relationships between a classifier and a set of labels in which some positive genes are mislabeled as negative. In Figure A.1a, mislabeling is random with respect to classifier accuracy. In Figure A.1b, the classifier has lower specificity on mislabeled positives than on correctly labeled positives. In Figure A.1c, the classifier has higher specificity on mislabeled positives than on correctly labeled positives, and in Figure A.1d, a mislabeled positive gene is less likely to be labeled positive than a negative gene.

In all cases except for the unusual scenario of Figure A.1d, estimated specificity underestimates true specificity. To see this, let 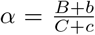 be the ratio of mislabeled negatives. If the classifier has any ability to discriminate mislabeled positives from negatives, then *B*/(*B* + *b*) < *C*/(*C* + *c*) and *B* < *αC*. Therefore,

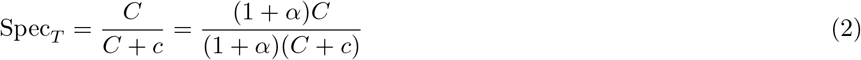

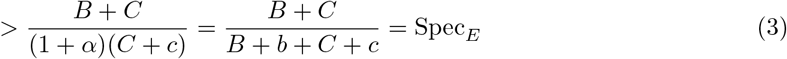

Therefore, in most cases, we can expect that specificity estimated from an imperfect GS gene set underestimaes true specificity.

However, sensitivity, may be biased either up or down. Sensitivity is only accurately estimated in the scenario illustrated in Figure A.1a. It will be over-estimated if the classifier is worse in the mislabeled positives than in the correctly labeled positives (scenarios in Figures A.1b and A.1d) and under-estimated if the classifier is better in the mislabeled positives (scenario in Figure A.1c).

### B Simulation Study

We use simulations to illustrate biased evaluation of data with a subset of mislabeled genes. In our simulations, each gene has two continuous features, P and E. We think of these as continuous measurements of the evidence that a gene affects a trait through either protein variation (P) or through expression variation (E). Let *Y*_*i*_ be a binary indicator that gene *i* is causal for the trait of interest. We consider two simulation scenarios. In both cases, *P*_*i*_ and *E*_*i*_ are generated from independent standard normal distributions and *Y*_*i*_ is generated as

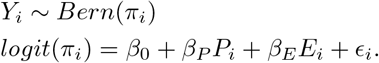

In Model 1, the “protein-dominant” model, *β*_0_ = *−*3, *β*_*P*_ = 6, and *β*_*E*_ = 2. In Model 2, the “expression dominant” model, *β*_0_ = −3, *β*_*P*_ = 2, and *β*_*E*_ = 6.

In each simulated data set, we generate *P, E*, and causal status, *Y* for 20,000 genes that are divided into a set of 10,000 genes used for training and 10,000 genes used for testing. In the training set, we fit two classifiers, the P-classifier, and the E-classifier, by fitting a logistic regression with *Y* as the outcome and either *P* or *E* as a predictor. Most causal gene discovery methods are not trained on a data set with known labels. However, this strategy gave us a straight-forward method to obtain classifiers that would be more sensitive for genes with high *P* or *E* values.

**Fig. A.1:**
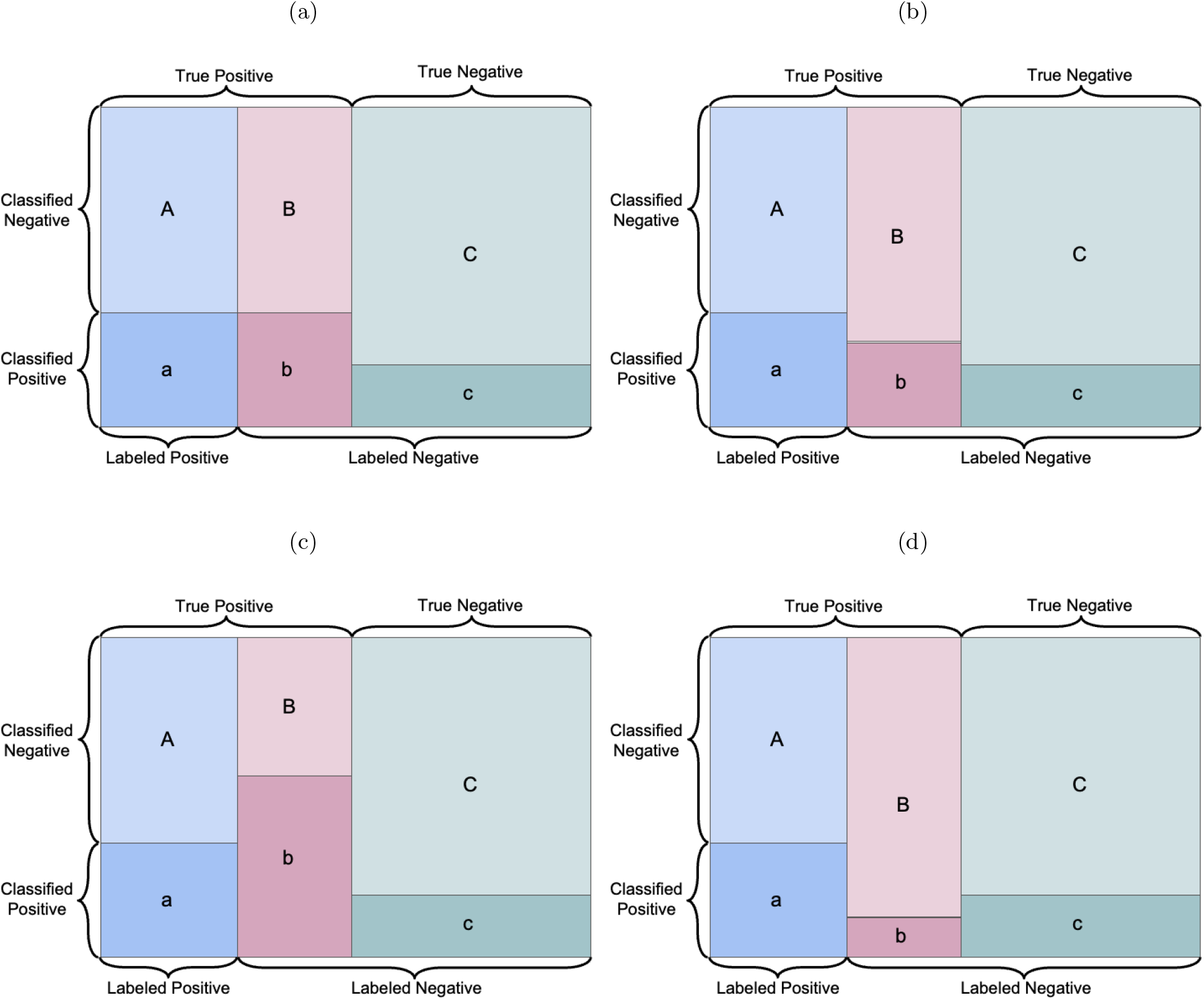
Possible relationships between classifier and imperfect labels. Each rectangle represents the total set of labeled genes. Columns indicate the three types of labeled genes, correctly labeled positive genes (blue, A/a), positive genes labeled as negatives (red, B/b), and correctly labeled negative genes (green C/c). Horizontal divisions indicate classifications, with the lower divisions (lowercase letters, darker shading) classified as positive and upper divisions (capital letters, lighter shading) classified as negative.

The 10,000 genes in the testing set function as our GS gene set. We consider 3 possibilities. Either all genes are correctly labeled, positives with high levels of *P* are more likely to be correctly labeled, or positives with high levels of *E* are more likely to be correctly labeled. We refer to these as correct labels, P-enriched labels, and E-enriched labels. Let *Z*_*C,i*_, *Z*_*P,i*_, and *Z*_*E,i*_ be correct, P-enriched, and E-enriched labels for gene *i* in the testing set. We generate these as *Z*_*C,i*_ = *Y*_*i*_, *Z*_*P,i*_ = *Y*_*i*_*W*_*P,i*_, and *Z*_*E,i*_ = *Y*_*i*_*W*_*E,i*_ with

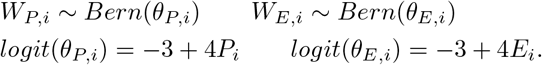

In Model 1, the P-enriched and E-enriched labels misclassify 4.56% and 13.5% of positive genes respectively. In Model 2, P-enriched labels misclassify 13.6% and E-enriched labels misclassify 4.46% of positive genes. ROC curves estimated for each label set under both models are shown in Figure B.2. Results presented in main-text Figure 1c correspond to Model 2 (larger E-feature effect) with P-enriched labels. In all cases, label enrichment results in biased estimation of model performance. When P-enriched labels are used, AUC of the P-classifier is over-estimated and AUC of the E-classifier is underestimated. This pattern is reversed using E-enriched labels. In two cases (Figures B.2b and B.2c) this results in a misordering of the two classifiers.

**Fig. B.2:**
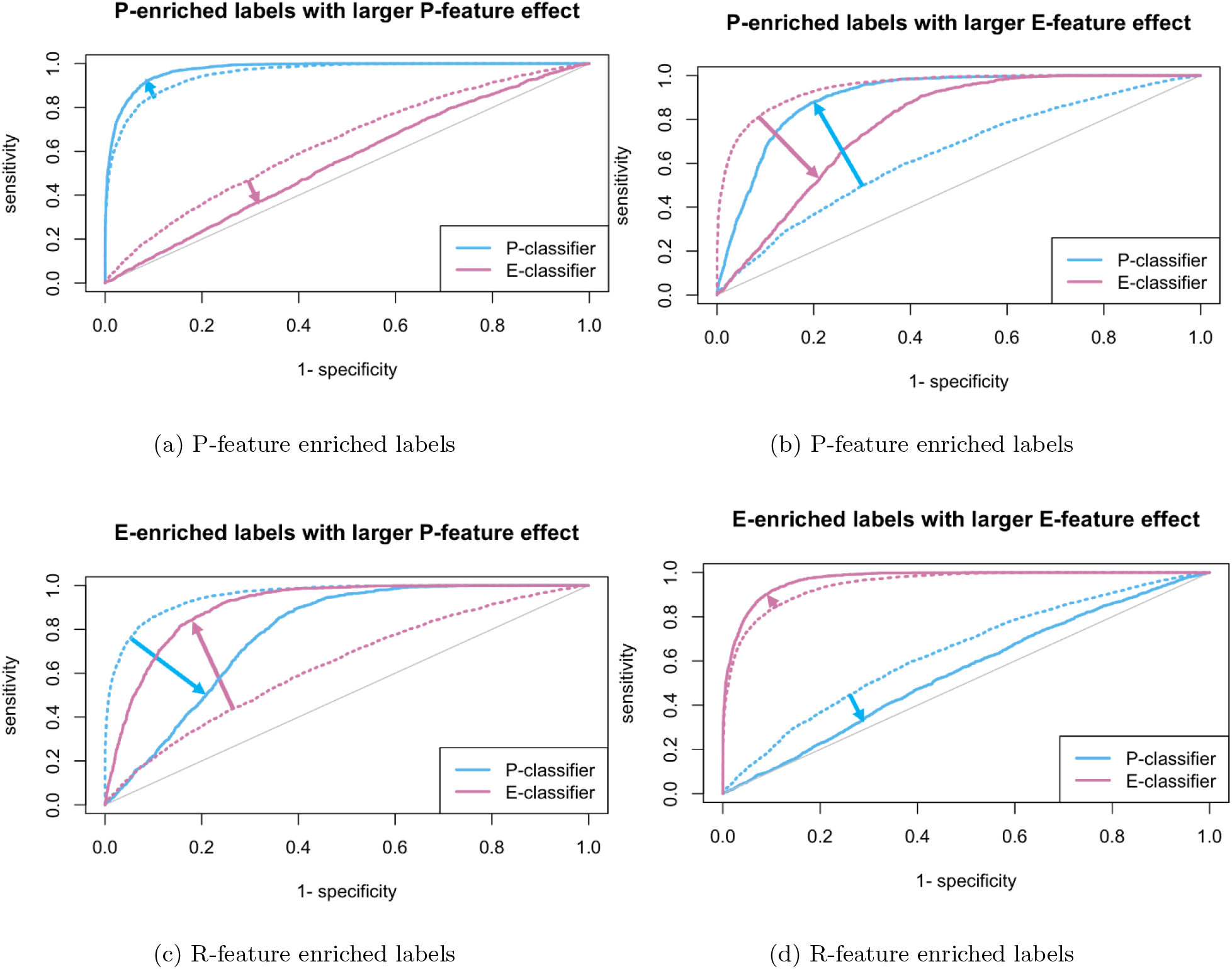
Comparison of Classifiers on True vs. Biased Labels. Classifiers evaluated against true labels (dotted lines) and biased labels (solid lines). Arrows highlight the change in ROC curve when some true positive labels are missed.

## Footnotes

1 This article adopts a PU scheme when using the GS gene set in benchmarking classification methods.

2 Causal genes selected here are not used to evaluate classification methods but to narrow down putative causal variants.

## Notes

https://lijiaw.gitlab.io/GS-gene-sets/comparison.html

### Competing Interest Statement

The authors have declared no competing interest.

### Summary of Updates

Clarified description of simulation construction procedure in Appendix B and updated the corresponding figures.

